# Fixational eye movements as active sensation for high visual acuity

**DOI:** 10.1101/2022.04.26.489583

**Authors:** Trang-Anh E. Nghiem, Oscar Dufour, Jenny L. Reiniger, Wolf M. Harmening, Rava Azeredo da Silveira

**Author notes:** All authors designed research. TAEN and RAdS designed the model and TAEN implemented it and ran simulations. JLR and WMH designed the experimental paradigm and conducted experiments. TAEN and OD analyzed experimental data. TAEN and RAdS wrote the manuscript and all authors edited it. The authors declare no competing interests.

## Abstract

Perception and action are inherently entangled: our world view is shaped by how we explore and navigate our environment through complex and variable self-motion. Even when fixating on a stable stimulus, our eyes undergo small, involuntary movements. Fixational eye movements (FEM) render a stable world jittery on our retinae, which contributes noise to neural coding. Yet, empirical evidence suggests that FEM help rather than harm human perception of fine detail. Here, we elucidate this paradox by uncovering under which conditions FEM improve or impair retinal coding and human acuity. We combine theory and experiment: model accuracy is directly compared to that of healthy human subjects in a visual acuity task. Acuity is modeled by applying an ideal Bayesian classifier to simulations of retinal spiking activity in the presence of FEM. In addition, empirical FEM are monitored using high-resolution eye-tracking by an adaptive optics scanning laser ophthalmoscope. While FEM introduce noise, they also effectively pre-process visual inputs to facilitate retinal information encoding. Based on an interplay of these mechanisms, our model predicts a relation between visual acuity, FEM amplitude, and single-trial stimulus size that quantitatively accounts for experimental observations and captures the beneficial effect of FEM. Moreover, we observe that human subjects’ FEM statistics vary with stimulus size, and our model suggests that changing eye motion amplitude, as the subjects indeed do, enhances acuity as compared to maintaining eye motion size constant. Overall, our findings indicate that perception benefits from action even at the fine and noise-dominated spatio-temporal scale of FEM.

**Significance Statement:** Perception is inherently active: we need to move our eyes to see the world around us. Yet our eyes also undergo tiny, unconscious movements that can blur out fine visual details. Paradoxically, previous work suggested that these small movements aid fine detail perception. Here, we investigate this paradox to uncover in which contexts small eye movements help or harm visual acuity. Comparing a model of retinal responses with recordings of human visual acuity, we elucidate the mechanisms by which and conditions in which small eye movements support fine detail discrimination. Our results also suggest that varying eye movement amplitude according to stimulus size enhances retinal coding, highlighting that perception is active even at the level of very fine eye movements.

---

**H**uman perception is inherently active: as you read this sentence, your eyes produce incessant, intricate movements necessary for vision. Even when attempting to fixate a single letter without moving, spontaneous, small-amplitude eye jittering occurs, known as fixational eye movements (FEM). Yet, in spite of FEM, we perceive our visual environment as stable (1) and we are able to distinguish details finer than the amplitude of FEM (2). Our understanding of the role of FEM in shaping perception remains incomplete: on the one hand, FEM can impair visual acuity by introducing noise (2, 3); on the other hand, experiments show that FEM can enhance fine detail vision in human subjects (4, 5).

As evidence suggests that information about the trajectories traced out by the eyes during FEM is unused in downstream processing (3, 6–9), one expects FEM to blur out fine visual details and, thereby, hinder perception. Indeed, retinal spiking is stochastic by nature, and therefore keeping one’s eyes static to overcome noisy spiking is advantageous compared to moving them. A static eye leads to more spikes encoding the same stimulus in the same position, which allows the noise to be averaged out over time. This effect, according to which FEM are harmful for coding, is illustrated in previous modeling works in which a Bayesian decoder infers stimuli pixel-by-pixel from simulations of stimulus-evoked retinal spiking (2, 3). In this framework, small to no FEM allow for a decoder to average out noise, as compared to larger FEM, though it remains possible to decode stimuli in the presence of sufficiently small FEM.

In a contrasting picture, human psychophysics suggest that FEM support perception: experimentally suppressing FEM by stabilizing stimuli on a subject’s retina impairs visual acuity (4), especially in distinguishing fine details (5). Here, FEM play a helpful role by pre-processing the visual input so as to effectively transform the spatio-temporal structure of receptive fields in retina (10–12) and thalamus (13, 14). In the temporal domain, FEM renders retinal responses to stimuli less transient in time: while retinal spiking decays over time upon presentation of a static stimulus, jittery eye motion refreshes each cell’s receptive field, which elicits sustained retinal activity that encodes the visual input (10). In the spatial (frequency) domain, the statistics of FEM are such that the pre-processing filter they effectively apply can enhance retinal gain at high spatial frequencies: small-amplitude jitter in eye position causes fine spatial stimulus features to move through retinal receptive fields, which induces temporal changes in receptive field contents and hence stimulus-relevant variations in retinal activity (11, 12). Overall, FEM allow for temporal encoding of stimuli by rendering retinal activity sustained through time and able to encode fine spatial detail through temporal variations in spiking.

Here, we combine theory and experiment to study the role of FEM in shaping retinal coding and visual acuity. In particular, we investigate the interplay of the opposite mechanisms of averaging over noisy spikes, which renders FEM harmful, and temporal coding, which makes FEM advantageous for visual acuity. The two mechanisms are most relevant at different stimulus sizes and FEM amplitudes; we therefore study their interactions and the resulting nature of visual coding across experimental conditions and model parameters.

From the interplay of different coding mechanisms, we derive a relation between visual acuity, FEM amplitude, and stimulus size, which we verify experimentally. In particular, our model accounts for FEM enhancing acuity for stimuli as small as receptive field size or smaller, consistent with previous reports (5, 9). Further, our experimental results show that subjects’ FEM amplitude varies depending on stimulus size. This has potential benefits for retinal coding, as modeling suggests that enhanced acuity can be achieved by varying FEM amplitude as subjects do. The results highlight that perception is inherently active even at the level of FEM.

## Results

To understand how FEM statistics affect visual acuity, we first study FEM trajectories during a visual discrimination task, in which 17 human subjects must discriminate between orientations of a Snellen E in a four-alternative forced choice task, where the letter E is displayed at adaptive stimulus sizes quantified by the spacing between adjacent bars of the “E” (Fig. 1a), between 0.3 and 0.8 arcmin. FEM trajectories are recorded through eye tracking with an adaptive optics scanning laser ophthalmoscope (AOSLO) (one subject’s example trajectories shown in Fig. 1b). The results reveal trajectories consistent with a random diffusion process, in that power scales as 1/*f*^2^ with frequency *f* (Fig. 1c), as expected from the literature (15), and the squared end-to-end length of trajectories grows linearly with time (Fig. 1d, see Supplementary Information for derivations in the case of a diffusion process). The slope of this proportionality relation, governed by the diffusion constant *D* in a diffusion process, varies substantially across subjects (Fig. 1e).

**Fig. 1.**
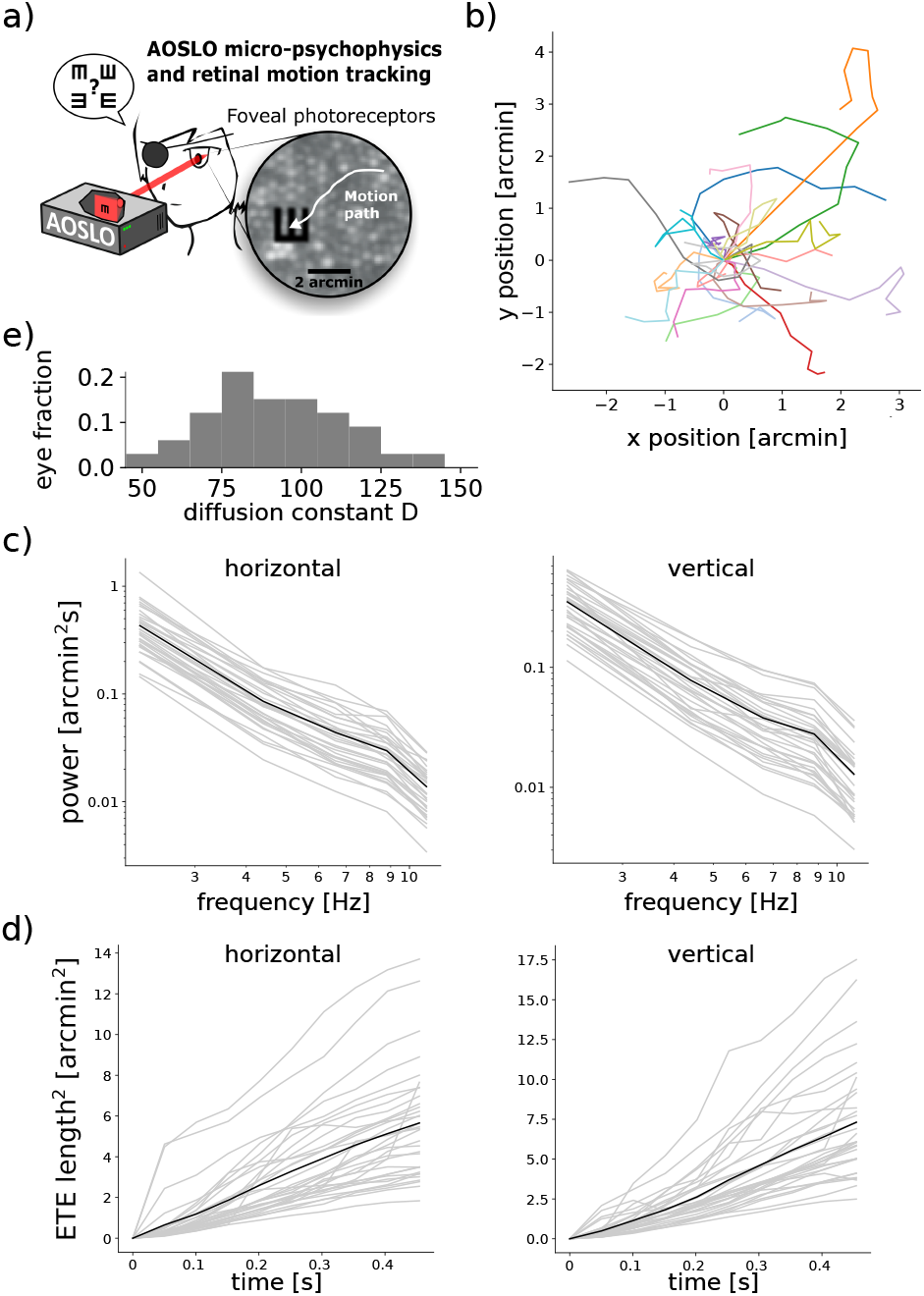
FEM are consistent with a random diffusion process, with diffusion constant varying importantly across subjects. (a) Experimental set-up and frame captured by the AOSLO, showing a subject’s retinal photoreceptor mosaic with a visual stimulus and recorded FEM trajectory. (b) Example FEM trajectories aligned to their starting point for one subject across stimulus presentations (colors). (c) Power spectra of FEM trajectories averaged over trials, shown for each subject eye (in grey, mean in black) along horizontal (left) and vertical (right) directions. (d) Mean square end-to-end distance of trajectories shown for each subject eye (in grey, mean in black) along horizontal (left) and vertical (right) axes. (e) Histogram of diffusion constants *D*, fitted over all trajectories of all trials and stimuli, for each subject. The results report important inter-subject variability in terms of *D*, with the largest *D* almost three times larger than the smallest value.

How do FEM affect visual discrimination? From a theoretical point of view, we must consider the interplay between two opposing coding mechanisms which involve the statistics of FEM, controlled by *D*, the stimulus properties, and the response properties of retinal cells (Fig. 2a). On the one hand, coding benefits from averaging over noisy spikes by keeping the eyes stable (2). Averaging over noise especially supports acuity in the case of coarse stimuli, where stimulus features are as wide as receptive fields or wider, and stimulus orientation can still be identified when the stimulus is downsampled into the space of retinal receptive fields (Fig. 2b).

**Fig. 2.**
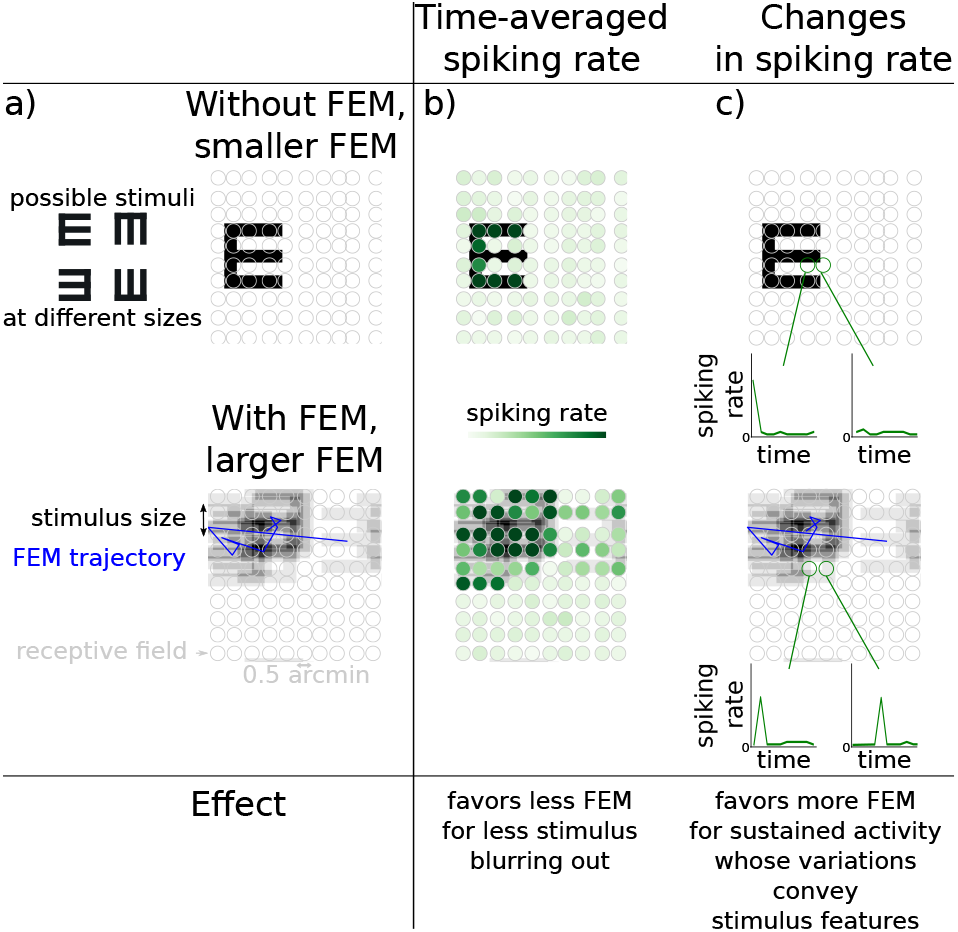
Mechanisms by which FEM affects retinal responses and stimulus encoding. (a) Snellen E stimulus (black) moving alongside schematic FEM trajectory (blue) and projected onto RGC receptive fields (grey) for stable stimuli (top) and jittery stimuli in the presence of FEM (bottom). Each letter E represents the input to the retina at a different point in time. (b) Time averaged spiking rate (green) elicited for each RGC by moving stimuli. In this coding scheme, large FEM result in a blurred out stimulus. Conversely, less to no FEM are favored as they allow to average out noise from stochastic spiking activity by keeping the eye relatively stable through time. (c) Changes in the spiking rate of two example RGCs due to variations in receptive field content caused by FEM. In the absence of FEM, light intensity is constant through time in each receptive field, leading to RGC activity decaying in time. In the presence of FEM, however, RGC responses to stimuli are sustained, and changes in spiking rate can convey stimulus-relevant information.

On the other hand, temporal encoding of visual stimuli by the dynamics of retinal activity benefits from FEM. Indeed, FEM refresh the content of receptive fields, which leads to more sustained stimulus encoding over time, as well as allow for FEM-induced temporal fluctuations in retinal responses, which can convey information about small stimulus details. For example, in Fig. 2d (bottom row), the stimulus moves in and out of a given receptive field under the effect of FEM, which results in changes in retinal spiking rate that encode relevant information about stimulus position and orientation. Temporal coding rendering FEM advantageous is consistent with FEM effectively transforming the structure of spatiotemporal receptive fields (10, 11), which is believed to enhance retinal sensitivity to fine stimulus details (5). This mechanism appears particularly relevant for stimulus features finer than receptive field size, for which retinal encoding amounts to downsampling, and intermediate FEM amplitudes. For small or vanishing FEM, receptive field contents are constant and retinal activity therefore decays over time (Fig. 2d top). Conversely, for FEM larger than the size of the small details, coding is impaired by the resulting uncertainty on stimulus location with respect to the eye. From the interplay between these coding mechanisms, one expects that visual acuity depends non-trivially on both stimulus size and FEM amplitude.

To explore the combined effect of these mechanisms, we investigate a simple model of retinal responses in the presence of FEM. We model FEM as resulting from a two-dimensional diffusion process with Poisson step size, controlled by a diffusion constant *D* (see Methods for details). We model the spiking activity of retinal ganglion cells (RGC) at the foveola, assuming one-to-one correspondence with photoreceptors (16), in response to stimuli moving in the retina’s reference frame. RGC receptive fields are arranged into a mosaic (Fig. 1a). Each RGC’s spiking rate in response to stimuli is characterized by a spatio-temporal kernel: spatial receptive fields are Gaussian with circular symmetry, and temporal receptive fields render RGCs sensitive to temporal changes in light intensity, as in transient RGCs. From simulated spiking rates, RGC spikes are drawn according to a Poisson process. The only free parameters are the temporal kernel time scale as well as the gain of RGCs and baseline level of RGC activity, which control the sensitivity to stimuli and the spiking rate in the absence of stimuli, respectively. The parameters are set based on the previous literature on RGC models (2, 3) informed by recordings in the primate retina (17, 18), in particular to ensure that RGC spiking rates remain lower than 200 Hz. Hence, the likelihood of spiking given the stimulus can be estimated precisely in the model.

The amount of stimulus-relevant evidence conveyed by RGC spikes can then be quantified across stimulus sizes and *D* values, with a Bayesian classifier as an ideal model for how the rest of the brain processes information from retinal spikes. This classifier is optimal in that it accumulates all the information available in spiking activity, thus yielding the best possible accuracy at classifying stimuli given the simulated retinal spikes. The classifier has access to the form of the likelihood of spiking given the stimulus, as well as to the statistics of the eye movements, i.e., *D*, but not their specific trajectory. Starting with no prior knowledge about the stimulus and hence a flat prior distribution, evidence is accumulated from RGC spiking patterns, and the posterior distribution is updated as follows:

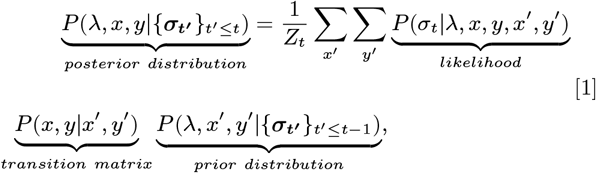

where *λ* denotes the stimulus orientation (top, bottom, left, or right), *x* and *y* are its centre position in the retina reference frame, {***σ_t’_***}_*t*’_ ≤ _*t*_ is the history of spiking patterns across RGCs up to time *t*, and *Z_t_* is a normalization constant. *P*(*σ_t_*|*λ,x,y,x*’, *y*’) is a product of 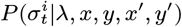 over all RGCs *i* (see Supplementary Information) since the cells are independent and any correlation between their activities comes from the stimulus. *P*(*x,y*|*x’,y’*) is the transition matrix governing the diffusion process of FEM, characterized by the diffusion constant *D* (see Supplementary Information); namely, the probability that the stimulus moves, during one time step, from coordinates (*x’,y’*) to coordinates (*x,y*).

The orientation of the E for which the posterior is maximized is the classifier’s outcome: *λ*\* = argmax_*λ*_Σ_*x*_Σ_*y*_*P*({***σ*_*t*’_**}_*t*’≤*t*_|*λ,x,y*). Repeating this procedure for 50 trials, one can estimate a fraction of correct discrimination after 500 ms of stimulus presentation (to match experimental conditions), for each value of *D* and each stimulus size.

Inspecting how the model gathers evidence from spikes to estimate stimulus orientation, we find that learning occurs for a large range of values of *D* as the fraction of correct discrimination increases over time, albeit at different rates depending on *D* and stimulus size (Fig. 3). For large stimuli (Fig. 3a), all values of *D* allow for near-perfect accuracy, quantified by a fraction of correct discrimination nearing 1, except for *D* = 0; in this case, learning occurs slowly because cells spike sparsely. Intermediate values of *D* (between 100 and 300) yield enhanced fractions of correct discrimination. Finally, a larger value of *D* is associated with lower fraction of correct discrimination for stimulus sizes smaller than the receptive field size, as well as slower learning for larger stimulus sizes. This can be understood in terms of the interplay between mechanisms: very small *D* does not allow refreshing receptive fields or encoding through differences in spiking rate (Fig. 2d), while very large *D* prevents staying around the same position for long enough to average over spiking activity (Fig. 2c). The optimal value of *D* is observed to vary non-trivially with stimulus size due to interactions between the opposing coding mechanisms. Indeed, too small eye movements do not enable time-sustained coding with stimulus-relevant time fluctuations in RGC spiking rate, and too large values of *D* are detrimental to coding as averaging over noise is impaired when stimuli moves too fast with respect to the retina. Overall, the interaction between mechanisms gives rise to an optimal value of *D* that increases first and then decreases with stimulus size in model predictions, which can be tested in empirical data.

**Fig. 3.**
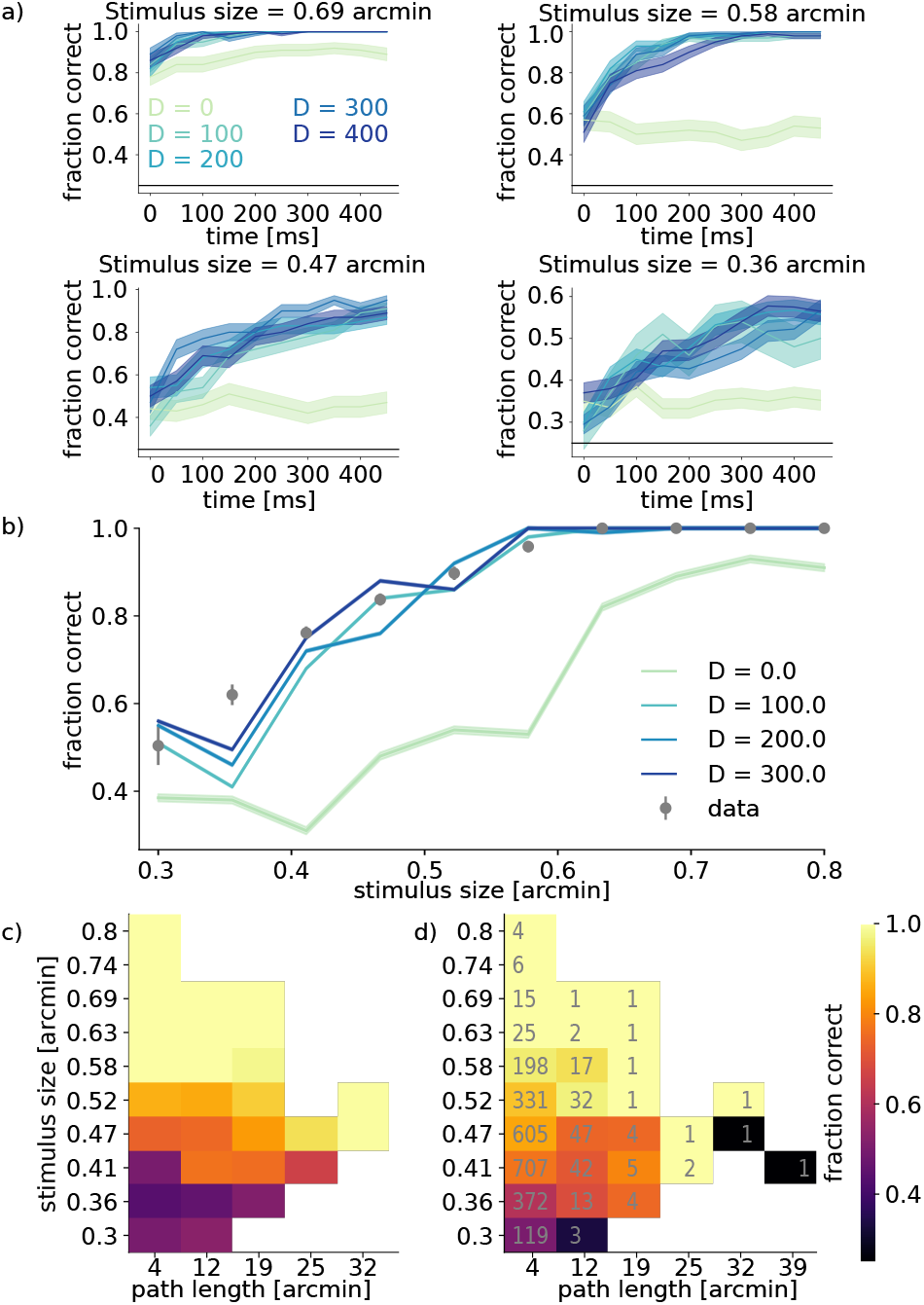
Bayesian classifier predicts that accuracy in visual discrimination tasks non-trivially depends on stimulus size and FEM amplitude, which is verified in empirical data on a trial-by-trial basis. (a) Classifier accuracy through time for different values of *D* (darker blue: larger *D*) and different stimulus sizes (decreasing size from top to bottom and left to right). Black line denotes the chance level fraction correct at 0,25. All fractions of correct discrimination increase through time. (b) Empirical (gray circles) and model-predicted (blue lines, darker blue for larger *D*) fraction of correct discrimination after 500 ms of simulation, as a function of stimulus size. Fraction of correct discrimination increases with stimulus size within a similar range for data and model, however the absence of FEM (*D* = 0) impairs accuracy across stimuli.(c-d) Heatmap of fraction of correct discrimination in bins of path length and stimulus size in the model (c) and data (d). For empirical values, the numbers in gray denote the number of trials per bin. Only bins where empirical path lengths were empirically recorded for those particular stimuli were represented. In these bins, the model predictions agree with empirical findings.

We compare the fraction of correct 500 ms trials from simulations and human subjects. First, we compute the mean fraction of correct discrimination over all subject eyes and trials, as a function of stimulus size. As expected, both the model and subjects discriminate large stimuli more accurately. Across values of *D*, the model displays good agreement with the data; as simulated fractions of correct discrimination across values of *D* lie in the same range as empirical ones, between 60% and 100% correct answers (Fig. 3b). It is also worth noting that the absence of FEM (*D* = 0, lightest teal line in Fig. 3b) yields poorer results, in agreement with empirical evidence that stabilizing stimuli on the retina impairs the ability to discriminate between stimuli (4, 5).

To investigate for which stimuli FEM are the most helpful, we examine the fraction of correct discrimination of the subjects as a function of both stimulus size and path length of FEM, defined as the total amplitude of the stimulus trajectory with respect to the eye in a trial. In order to compare with empirical observations, we consider the simulated fraction of correct discrimination for a range of values of *D* matching the range found in subjects (Fig. 1e). Then, we compute a weighted average of the fraction of correct discrimination across values of *D*, where weights correspond to how heavily represented each value of *D* is across subjects (Fig. 1e). Once again, we find agreement between data and model (Fig. 3c,d). For stimuli larger than the typical size of RGC receptive fields at the preferred retinal location of fixation on the retina (0.5 arcmin), discrimination is near-perfect across path lengths. However, for finer stimuli, our results suggest that the fraction of correct discrimination improves for larger path length, down to very fine stimuli at the limit of human acuity (0.3 arcmin).

Which mechanism explains the beneficial effect of FEM for visual decoding and acuity? The more immediate answer, namely that FEM refresh the image on the retina and thereby enhance spiking, is insufficient. To illustrate this point, we repeated our theoretical study using modified model RGC with a monomodal temporal filter, such that the model RGC responds to light intensity rather than temporal variations of it (*w* = 0 in Eq. 4). In this case, RGC response is not transient, and FEM are no longer necessary to elicit sustained RGC spiking. In this modified model, overall acuity deteriorates as expected, which confirms the relevance of the transient nature of RGC spiking to reproduce experimental results. Still, however, larger FEM path length boosts the fraction of correct discrimination for stimuli as fine as or finer than the receptive field size, consistent with empirical data (Supplementary Information, Fig. S1). Thus, the ‘refresh mechanism’ is not alone responsible for the benefits of FEM; in addition, temporal modulations of RGC activity caused by FEM convey stimulus-relevant information that enhances the visual acuity of an ideal decoder.

Surprisingly, we also note that the range of observed path lengths depends considerably on stimulus size. Indeed, path lengths are found to be up to five times larger for intermediate stimulus size (0.5 arcmin, comparable to RF size) than for the largest stimulus size (0.8 arcmin).

In sum, our results show that the empirical and model amplitude of FEM influences visual coding and discrimination on a trial-by-trial basis. Especially for stimuli finer than receptive fields, for which coding through changes in neural activity upon eye motion is especially relevant, longer FEM trajectories are favored. The model quantitatively captures experimental results by taking this mechanistic element into account. Indeed, our model explains how fine detail vision deteriorates in the absence of FEM, consistent with the experimental literature (5). Additionally, simulations quantitatively reproduce variations of the fraction of correct discrimination as a function of stimulus size and path length of FEM. Note that among model parameters, only *D* is fitted from empirically recorded FEM trajectories, and no parameters are adjusted to reproduce subjects’ fractions of correct discrimination. Surprisingly, the path length empirical FEM is find to vary depending on stimulus size - are these variations robust, and if so, do they influence retinal coding?

Quantifying the variations of FEM path lengths with stimulus size, we observe that significant differences can be noted overall in path lengths across stimulus sizes (Fig. 4a, Kruskal-Wallis test, *H* = 99, *p* = 10^-17^). In particular, statistical testing shows that path lengths for stimuli between 0.4 arcmin and 0.5 arcmin, near the subjects’ acuity threshold, were significantly different from others. Can observed changes in FEM amplitude across stimuli support retinal coding and visual acuity? We examine whether varying FEM amplitude across stimulus sizes, as subjects do, leads to improved acuity, as compared to keeping FEM amplitude fixed or varying FEM amplitude in different ways.

**Fig. 4.**
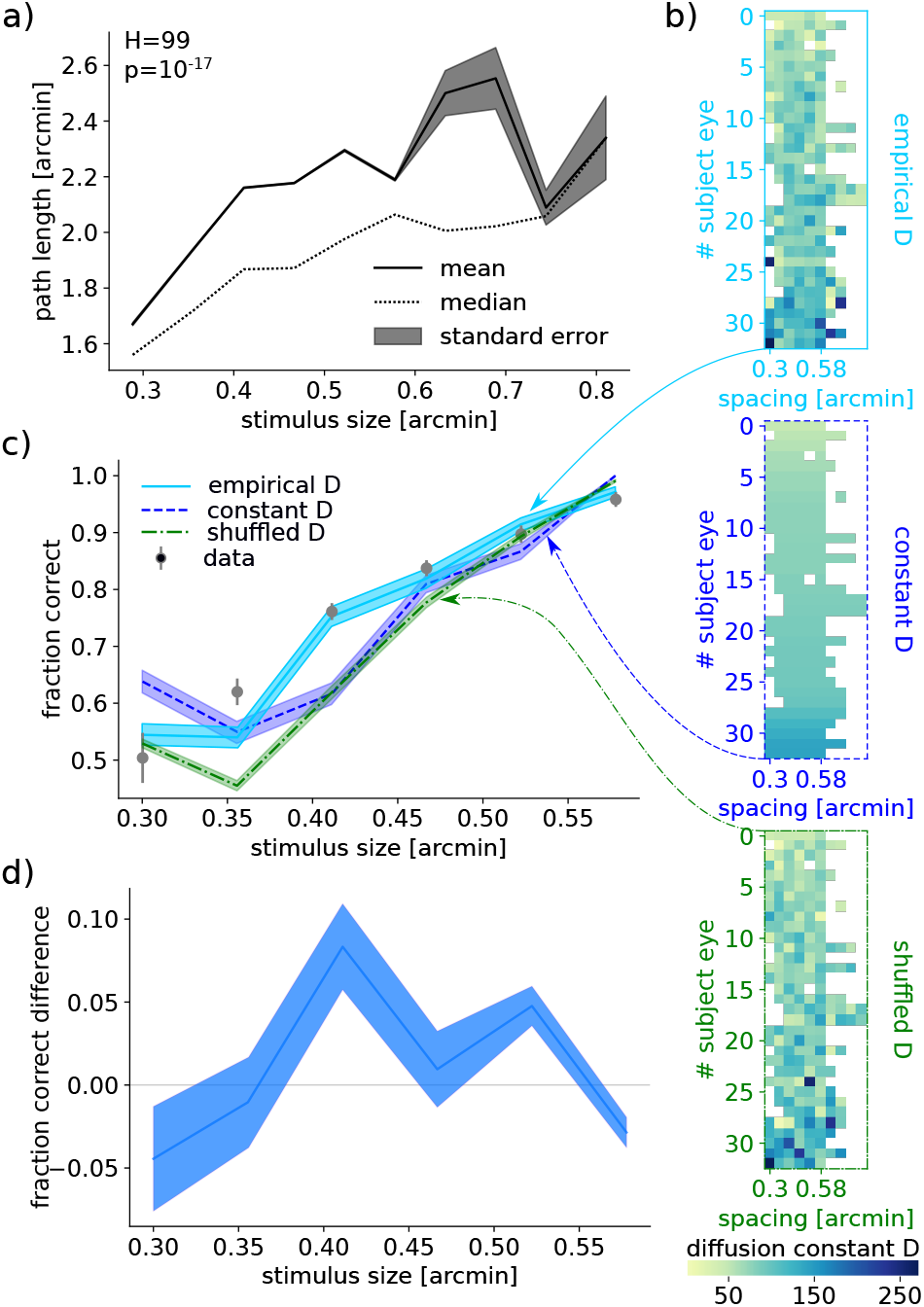
Variations in subjects’ FEM amplitude depending on stimuli lead to enhanced retinal coding. (a) Path length statistics as a function of stimulus size: the mean (black line), standard error of the mean (shaded area), and median (dotted line) are computed over all FEM trajectories for all subject eyes within bins of stimulus size. (b) Model-predicted fraction of correct discrimination for empirical stimulus-dependent (top), stimulus-independent (middle) *D*, and shuffled *D* across stimuli (bottom) shown as function of stimulus size. (c) Empirical (gray circles) and model-predicted fraction of correct discrimination for empirical (solid line), constant (dashed), and shuffled (dash-dotted) D. (d) Model-predicted difference in fraction of correct discrimination between trials with empirical *D* varying across stimuli (b, solid) and fixed *D* chosen to be the mean over sizes (b, dashed), as a function of stimulus size. Lines represent means, and shaded areas represent standard errors in the mean.

To that purpose (Fig. 4), we fit *D* separately for each stimulus size and each subject (Fig. 4b top); we compare this case to that of a fixed value of *D* equal to the average empirical *D* over stimuli (Fig. 4b middle), and to that of values of *D* shuffled across stimulus sizes (Fig. 4b bottom). Using our model, five trials are then simulated for each value of *D* per subject eye and per stimulus size, successively for empirical, averaged, and shuffled values of *D*. From the fraction of correct discrimination (Fig. 4c), we observe that varying *D* across stimuli according to empirical values yields an accuracy at the task similar to that of human subjects, as well as consistently enhanced acuity over keeping *D* constant or using shuffled values of *D* (Fig. 4d).

We note that free parameters in the model are not readjusted to resemble psychophysical results; as they are chosen to reflect RGC electrophysiological properties and kept constant throughout the manuscript. In particular, we observe that for a certain range of stimulus sizes smaller than foveal RGC receptive fields and near the limit of human acuity, the fraction of correct discrimination is enhanced on average by over a standard error in the mean and up to 10% when modeling empirical, stimulus-dependent *D* compared to when modeling averaged, stimulus-independent *D* or values of *D* shuffled across stimulus sizes (Fig. 4c). For smaller stimulus sizes (0.3 arcmin), the subjects and model perform poorly regardless of the values assigned to *D*. For stimuli larger than typical receptive field size (0.5 arcmin), the subjects and model perform near-perfectly across all values of *D*.

We conduct several additional analyses (data not shown) to check the robustness of our results. For example, we apply different shuffling protocols, that conserve the dependence of *D* on either fine or coarse scales of stimulus size; these analyses yield similar performance. We cannot exclude that the form of variations in *D* is incidental to this specific behavioral experiment, and happens to yield a superior performance in the model. Indeed, we are not able to characterize a simple form of the variations of *D* as a function of stimulus size. Yet, the conclusion that a variable value of *D* enhances visual acuity remains. The variations in *D* measured in human subjects suggest that, not only can FEM benefit the resolution of fine visual details, but they can do so actively through modulations of their amplitude according to stimulus size. Thus, vision may be active even at the scale of FEM.

## Discussion

In this work, we investigated the conditions under which FEM help or harm retinal coding and visual acuity. Empirical FEM trajectories were recorded in healthy human subjects during a discrimination task, and empirical FEM amplitudes were estimated for all subjects and stimulus sizes (Fig. 1). FEM amplitude and its relation to stimulus size non-trivially affect acuity, through an intricate interplay of different mechanisms: averaging over noisy spiking RGC activity, and temporal coding through refreshing the content of receptive fields, resulting in stimulus-informative variations in spiking (Fig. 2). Using the output of a model of retinal response to diffusing visual stimuli, an ideal Bayesian classifier successfully accounted for the benefits of FEM to discriminate fine stimuli (Fig. 3). The model of fraction of correct discrimination was found to qualitatively match human subject performance at the task, even though no model parameters were fitted to empirical fractions of correct discrimination, and only the diffusion constant of modeled FEM was fitted to recorded FEM trajectories. In addition, we noticed that subjects’ FEM path lengths varied with stimulus size. Our model predicted that empirically observed changes in FEM size across stimuli lead to improvements in visual acuity (over fixed-sized FEM), suggesting that perception can benefit from being active even at the scale of FEM (Fig. 4).

While earlier modeling work on interpreting spikes in the presence of FEM had suggested stimulus encoding and decoding was possible in spite of FEM (2, 3), our work establishes that FEM are able to not only allow, but also to enhance the encoding and decoding of task-relevant information, thereby accounting for experimental observations (5). Furthermore, our work uncovers a relation between the (fine) spatial scale of visual stimuli and the role of the FEM amplitude. When one considers the opposing mechanisms of noise averaging (which favors lower values of *D*) and of the response dynamics (which favors higher values of *D*), non-zero, optimal value of *D*, varying with stimulus size, emerges. This notion of optimal FEM statistics for stimulus encoding is complementary with existing ideas that focused on stimulus statistics, suggesting that scale-invariant eye movements effectively transform the visual input so as to enhance our perception of fine detail (5, 10). Our results quantitatively addresses the statistics of FEM and shows that optimal values of *D* for the encoding stimulus-relevant information vary depending on spatial frequencies present in stimuli. Recent modeling work exploring yet an additional mechanism, namely that encoding can take advantage of spatial heterogeneities in the retinal receptive field mosaic through FEM, also found benefits of FEM to visual coding (19). Our conclusions are complementary in that they exploit different mechanisms through which FEM enhance retinal coding.

Our experimental findings indicate that subjects can change the amplitude of their FEM during sustained fixation depending on stimulus size. This agrees with observations in the recent literature that empirical FEM statistics change in a task-dependent manner. In particular, the amplitude, speed, and curvature of FEM drift and microsaccades were reported to be distinct between passive viewing and acuity tasks (20). In addition, recent work in primates supports that FEM originate from central neural circuitry generating noise that controls FEM statistics (21), which is consistent with subjects’ ability to modulate FEM amplitude according to stimuli.

While we have discussed coding mechanisms at the retinal level, other mechanisms and their associated costs may also incur, including motor costs in quelling noisy and jittery motion to reduce FEM amplitude. There may be conditions under which FEM amplitude is suboptimal, and acuity is enhanced by partially stabilizing stimuli on the retina (22). The perspective we propose on the effect of FEM on perception provides quantitative predictions for how subject acuity varies with FEM amplitude, to be tested in future psychophysical experiments in which FEM size may be effectively enhanced or reduced by amplifying or compensating for motion through eye tracking. More broadly, our findings suggest that active sensing is relevant even down to fine spatio-temporal scales where noise plays a key role in shaping neural coding and sculpting human behavior.

## Materials and Methods

### Adaptive optics microstimulation

High resolution retina tracking during presentation of a small optotype was achieved by employing a custom adaptive optics scanning laser ophthalmoscope (AOSLO). In such a system, cone-resolved imaging and presentation of a visual stimulus is accomplished concurrently (23–25) (see SI for details).

### Visual acuity task and retina tracking

All 17 adult participants (8 male, 9 female, age: 10 – 42 years) took part in a visual acuity task. Participants had to indicate the orientation of a Snellen-E stimulus, presented randomly at one of four orientations (up, down, left, right). Each stimuli was presented for 500 ms at the center of the AOSLO raster after the participants initialized a trial with a keyboard press. Stimulus size during a trial was fixed, and was changed between trials following a Bayesian adaptive staircase procedure (QUEST (26)). Stimulus size decreased if the most recent response was correct, and increased at incorrect responses. Each experimental run consisted of 20 trials yielding stimulus gap sizes roughly between 0.3 and 0.8 arcmin. More than 20 trials were conducted in case some trials had to be discarded due to eye blinks, saccades, or eye tracking artefacts. Experimental runs were repeated five times for each eye of each participant (except one participant in whom only one eye was tested). With each stimulus presentation, a 1 s AOSLO video was recorded, from which retinal motion and stimulus position were extracted (27, 28). Retinal motion and thus eye motion traces during stimulation were extracted by a real-time, strip-wise image registration and stabilisation technique (29), with effective temporal sampling of ~ 960 Hz. Trials containing defects of eye-tracking stabilisation or microsaccades were identified by visual inspection (see SI for details).

### Retinal response model

Modeled RGC receptive fields are arranged into a mosaic reminiscent of Fig. 1a, which we simulate as a square lattice with jittered centre positions and receptive field sizes. Each receptive field is characterized by a spatial and temporal kernel. The spatial kernel is Gaussian with circular symmetry. The temporal kernel accounts for the fact that spiking activity decays to silence when the same fixed stimulus is shown within a time scale of the order of *dt* = 50ms (2), and is implemented by subtracting the light intensity observed 50ms ago, weighted by a kernel weight, from the current intensity within the receptive field. Subsequently, to obtain RGC spiking rates, we multiply the stimulus after applying the spatiotemporal receptive field kernel by a constant gain, and add a baseline that controls RGC spiking rate in obscurity. Then, spikes are drawn from a Poisson process with said spiking rate. Equation 4 describes RGC spiking rate and spiking probability in response to light stimuli.

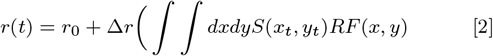

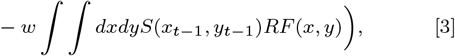

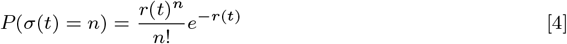

with *r*(*t*) the RGC’s spiking rate at time *t*, *r*_0_ = 100 Hz its baseline spiking rate in the absence of stimuli, Δ*r* = 500 Hz its gain controlling sensitivity to light intensity, *RF* its spatial receptive field specific to each neuron, 0 < *w* < 1 the temporal kernel weight, *S*(*x_t_, y_t_*) the two-dimensional stimulus centred onto position (*x_t_, y_t_*) the coordinates of the stimulus at time *t*, *σ*(*t*) the number of spikes fired by the RGC at time *t*, and *n* a non-negative integer. For computation purposes, all possible positions (*x,y*) are described by coordinates upon a discrete square grid and cyclic boundary conditions are applied. Parameter values for *dt*, *r*_0_, and Δ*r* are chosen based on retinal recordings (17, 18) and previous modelling work (2, 3).

## ACKNOWLEDGMENTS

The authors acknowledge Simone Blanco Malerba and Luc Stebens for useful discussion. Support for this research was provided by the CNRS through Unité Mixte de Recherche (UMR) 8023, the Swiss National Science Foundation Sinergia Project (CRSII5 173728), the Carl Zeiss Foundation (WMH: HC-AOSLO) and the Emmy Noether Program of the German Research Foundation (DFG) (WMH: Ha 5323/5-1).

## Supplementary Information for

### Supporting Information Text

#### Adaptive optics microstimulation

High resolution retina tracking during presentation of a small optotype was achieved by employing a custom adaptive optics scanning laser ophthalmoscope (AOSLO). In such a system, cone-resolved imaging and presentation of a visual stimulus is accomplished concurrently. The techniques have been described earlier (1–3), we mention only pertinent details here. In brief, the output of a supercontinuum light source (SuperK Extreme, NKT Photonics, Denmark) was spectrally filtered to create a red visible light channel, used for imaging, ocular wavefront sensing and microstimulation (center wavelength = 788 ±12 nm, FF01-788/12-25, Semrock, Rochester, USA). Adaptive optics correction, run in closed loop operation at about 25 Hz, consisted of a Shack-Hartmann wavefront sensor (SHSCam AR-S-150-GE, Optocraft GmbH, Erlangen, Germany) and a magnetic 97-actuator deformable mirror (DM97-08, ALPAO, Montbonnot-Saint-Martin, France). The imaging/stimulation beam was point-scanned across the retina, spanning a square field of 0.85 x 0.85 degrees of visual angle. The light reflected from the retina was detected in a photomultiplier tube (H7422-50, Hamamatsu Photonics, Hamamatsu, Japan) which was placed behind a confocal pinhole (pinhole diameter = 20 *μm*, equaling 0.5 Airy disk diameters). PMT signals were sampled at 20 MHz by a FPGA board (ML506, Xilinx, San Jose, USA), producing digital video frames at ~ 30 Hz with a spatial resolution of 600 pixels per degree of visual angle. By modulating the intensity of the imaging beam by an acousto-optic modulator (TEM-250-50-10-2FP, Brimrose, Maryland, USA), visual stimuli were created (thus a ‘light off’ stimulus within the visible scanning background, see main paper Fig. 1a).

#### Data pre-processing

Trials containing defects of eye-tracking stabilisation or microsaccades were identified by visual inspection of stabilized AOSLO videos and eye motion trajectories obtained from eye-tracking. Trials presenting sheared or trapezoid video frames associated with trajectories displaying large displacements within a single time frame were identified as stabilization failures and were discarded from subsequent analyses. For analysis, eye motion trajectories were downsampled to a 50 ms time bin to average over the AOSLO scanning over the recording frame pixel per pixel over 33 ms cycles.

#### Diffusion process and end-to-end length

Let ***R*** be the vector connecting the initial position of the eye to its position after *N_t_* steps of duration Δ*t*. We can write 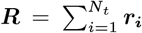, where the displacement vector at each of *N_t_* steps is given by ***r_i_*** = *l*(*X_i_**u_i_*** + *Y_i_**u_i_***); *X* and *Y* are iid Poisson random variables with mean *a* and variance equal to the mean, *ℓ* is the smallest possible non-zero step size, ***u_i_*** is a vector of norm 1 and phase drawn equiprobably from {0, *π*} and respectively ***u_i_*** is a vector of norm 1 and phase drawn equiprobably from {-*π*/2, *π*/2}. We compute the variance of the component of ***R*** along the x axis, 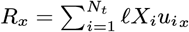.

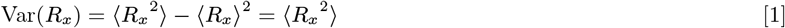

by symmetry of ***R_x_*** around 0 due to equiprobable leftward and rightward displacements.

The second moment can be calculated as follows:

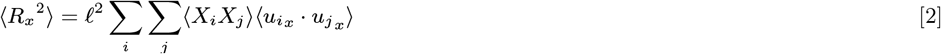

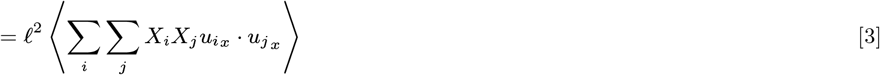

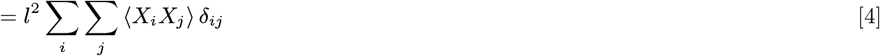

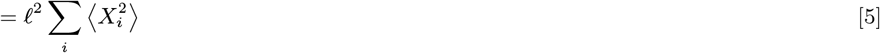

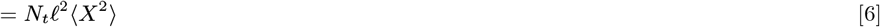

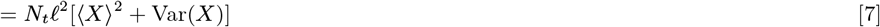

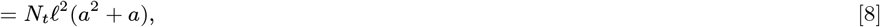

where we have used the properties of the Poisson process, *X*.

To compute *a* from 〈*R_x_*^2^〉, we solve the quadratic equation and retain the positive solution, 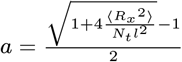. In addition, we recall that *a* = 2*D*Δ*t*. Therefore, by fitting the slope *α* of the square end-to-end length as a function of time, one can express the diffusion constant as a function of *α*, as

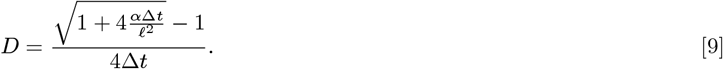

By symmetry, the same is true for the trajectory projection along the y axis.

#### Diffusion constant fitting

The diffusion constant can be estimated from the data by using Eq. 9, with *ℓ* =0.1 arcmin the minimum step size we used in discretizing FEM and *α* obtained by fitting the slope of the square end-to-end length in either x or y axe as a function of time. We note that if FEM were realizations of perfect random walks, with each step’s direction uncorrelated from the previous one, the same *D* value would be obtained by fitting from either FEM trajectory path lengths or end-to-end lengths. However, small discrepancies in *D* values may arise. We approximate FEM as a random walk in the interest of maintaining reasonable computational burden for Bayesian classification, even though evidence suggests that small directional correlations exist at longer time scales than our 50 ms time step, rendering FEM trajectories slightly smoother than those described by our simple random walk model (4, 5).

#### Power-spectral density

We want to compute the power-spectral density, *S*(*f*), of the random iid process, *R_x_* (and equivalently *R_y_*). By the central limit theorem, as *N_t_* tends to infinity, *R_x_* is well approximated by a Gaussian continuous random variable *W*(*t*), of mean 0 and variance 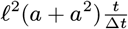 (Eq. 8, with *t* = *N_t_*Δ*t*). In the continuous limit, we have:

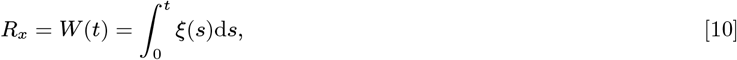

where *ξ*(*s*) is continuous white noise whose moments are 〈*ξ*(*s*)〉 = 0 and 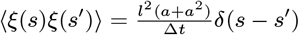. The power-spectral density is obtained as (6, 7):

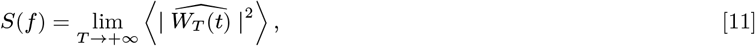

where 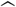 denotes the Fourier transform and *W_T_*(*t*) is *W*(*t*) truncated at time *T*. Explicitly, 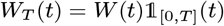 where 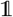 is the indicator function of the interval [0, *T*]. We use the following convention for the Fourier transform:

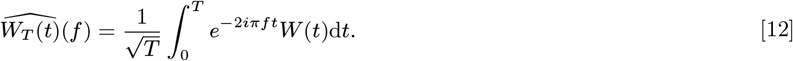

*S*(*f*) can then be written as

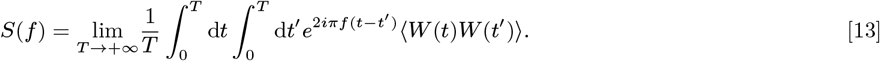

From Eq. 10, we can compute, the correlation term: 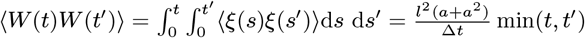. To compute *S*(*f*), first we divide the double integral on the square into an integral on the upper triangle and another on the lower triangle, to deal with the term min(*t, t*’). Then, recognizing that one double integral is the conjugate of the other, we rewrite the sum as twice the real part. The double integral is then calculated as:

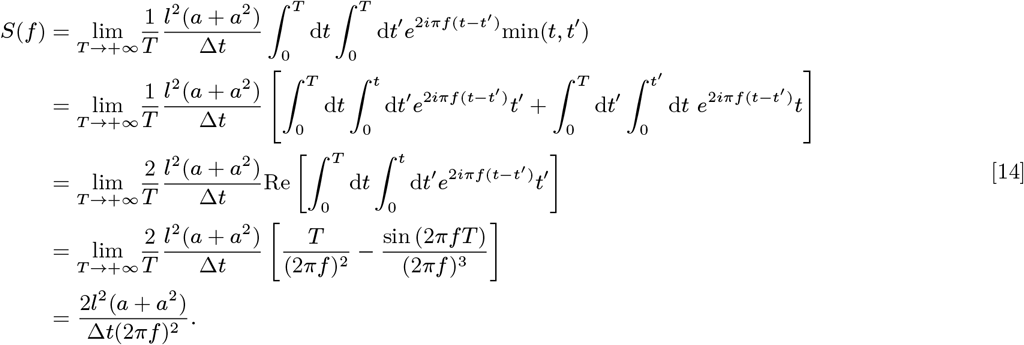

The power spectrum is inversely proportional to the square of the frequency. By symmetry, the same is true for the trajectory projected along the y axis.

#### Bayesian classifier

The classifier aims to output the stimulus orientation, *λ*, by accumulating evidence from RGC spiking over time. Initially, the prior distribution is flat, i.e., 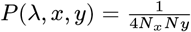 for each of the 4 possible stimulus orientations (top, bottom, left or right), each of the *N_x_* possible positions along the x axis and each of the *N_y_* possible positions along the y axis. Since the temporal kernel is non-vanishing over two time steps and the diffusion process is iid, the current position of the stimulus depends only on the position at the previous time step, and spikes from the current time step only carry information about stimulus position during the current and previous time step. The posterior distribution can therefore be updated from the following:

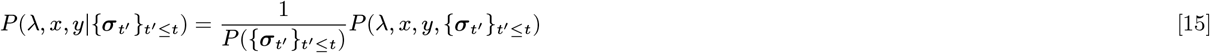

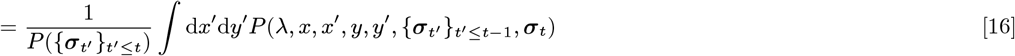

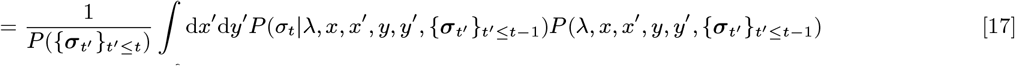

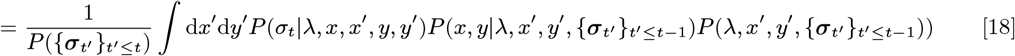

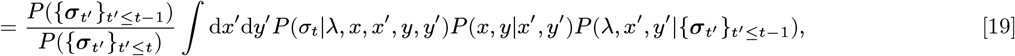

hence Eq. 1 of the main text. The transition matrix, *P*(*x,y*|*x*’, *y*’), containing the probabilities of all possible *x*’, *y*’ given the currently considered *x, y*, is written as

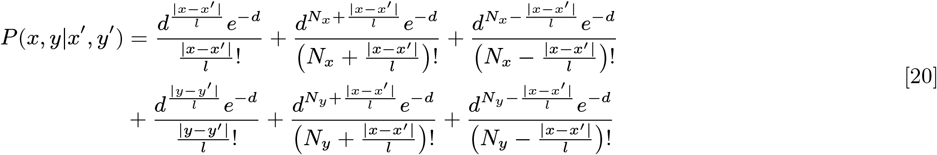

where *d* = 2*D*Δ*t*. The first three terms of the sum account for displacements along the x axis, where the first term captures the contribution of direct jumps from *x*’, *y*’ to *x, y*, the second term of longer jumps around the grid through the left, and the third term of longer jumps around the grid through the right due to cyclic boundary conditions. Similarly, the last three terms describe contributions from displacements along the y axis, including directly, around the grid through the top and the bottom.

---

**Fig. S1.**
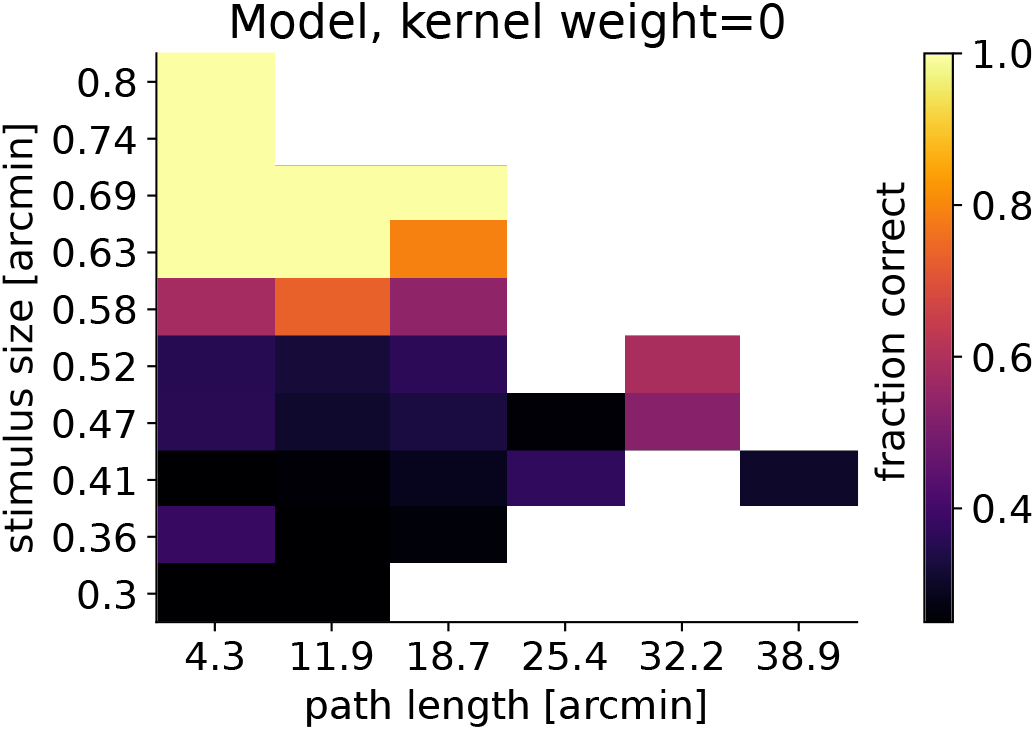
Model fraction of correct trials as a function of stimulus size and path length for sustained cells. Although acuity is impaired for finer stimuli than receptive field size, longer FEM trajectories still leads to improved fractions of correct trials compared to shorter trajectories.

